# Adipocyte lipolysis protects the host against *Trypanosoma brucei* infection

**DOI:** 10.1101/2022.11.05.515274

**Authors:** Henrique Machado, Peter Hofer, Rudolf Zechner, Luísa M. Figueiredo

## Abstract

*Trypanosoma brucei*, an etiological agent of African trypanosomiasis, colonizes the interstitial spaces of the adipose tissue in disproportionately high numbers, inducing a robust local immune response. Loss of body weight, including loss of adipose mass, is a hallmark symptom of African trypanosomiasis. Nevertheless, it is unclear which molecular mechanisms drive this loss of adipose mass and in turn whether it contributes to pathology. Here we show that lipolysis is activated early in infection in adipose tissue of *T. brucei-*infected mice. This activation is dependent on immune activation, as mice deficient for both B and T lymphocytes failed to upregulated adipocyte lipolysis upon infection and retained higher fat mass. Genetic ablation of the rate limiting adipose triglyceride lipase specifically from adipocytes in mice (*Adipoq*^*Cre/+*^*-Atgl*^*fl/fl*^) prevented the upregulation of adipocyte lipolysis during infection, leading to reduced loss of fat mass and adipocyte volume. Surprisingly infected *Adipoq*^*Cre/+*^*-Atgl*^*fl/fl*^ mice succumbed earlier to infection and presented a higher parasite burden in the gonadal adipose tissue, indicating that lipolysis limits the growth of the parasite population. Collectively, this work provides molecular mechanistic insight into the loss of fat mass in African trypanosomiasis and identifies adipocyte lipolysis as a host-protective mechanism during a *T. brucei* infection.

## Introduction

*Trypanosoma brucei* is an unicellular and extracellular parasite transmitted to humans and cattle through the bite of infected tsetse flies, causing Human and animal African trypanosomiasis (HAT and AAT), respectively^1^. Within the mammalian host, *T. brucei* colonizes both intravascular and extravascular spaces, causing a progressive wasting disease that is lethal if left untreated^2^.

The pathology of African trypanosomiasis is extensively modulated by the extravascular location of the parasite. This is well exemplified by the classical neurological symptoms observed in humans upon colonization of the central nervous system^3^ and, more recently, the skin lesions observed in HAT patients have been associated with the presence of *T. brucei* in the dermis^4^.

The loss of fat/adipose mass, which occurs during *T. brucei* infections in the context of a broader wasting syndrome^3^, may be associated with the parasite’s presence in the adipose tissue (AT). This possibility is supported by the fact that *T. brucei* accumulates in disproportionately high numbers in the AT of infected mice^5^. Moreover, in the gonadal AT (i.e. largest visceral AT depot), this colonization leads to a progressive accumulation of myeloid cells as well as tumor necrosis factor-alpha (TNF-α) and interferon-gamma (IFN-γ)-producing lymphocytes^6^. Concomitant to the mounting of this immune response, AT depots undergo a progressive loss of weight^5,6^, suggesting that AT colonization by *T. brucei* and the ensuing immune response may be responsible for the loss of fat mass observed in African trypanosomiasis.

Loss of AT depot mass occurs when lipid catabolism is favored over lipid anabolism (i.e. lipogenesis), resulting in mobilization of acyl-glycerol species contained within the adipocyte’s lipid droplet^7^. In adipocytes this mobilization occurs primarily through neutral adipocyte lipolysis, where triacylglycerol (TAG) is sequentially hydrolyzed into diacylglycerol and monoacylglycerol by three lipases: adipose triglyceride lipase (ATGL), hormone-sensitive lipase (HSL) and monoacylglycerol lipase^7^. Upon hydrolysis of acyl-glycerol species, the resulting free fatty acids and glycerol molecules are exported to the AT’s interstitial spaces and subsequently into the circulatory system.

A myriad of stimuli regulate adipocyte lipolysis, including inflammatory molecules (e.g. TNF-α)^8^, sympathetic nervous system cues (e.g. norepinephrine)^9^, hormones (e.g. insulin)^10^ and bacterial products (e.g. lipopolysaccharide and peptidoglycans)^11,12^. Proper regulation of adipocyte lipolysis is required for whole-organism energy homeostasis, especially during times of negative energy balance (e.g. fasting)^13^. Importantly, deregulation has been shown to promote the development of cancer-associated cachexia^14^, insulin resistance^15^ and hepatic steatosis^16^.

Here we show that loss of fat mass during infection occurs due to an increase in adipocyte lipolysis, particularly through ATGL activity. Moreover, we show that the immune response is an important driver of adipocyte lipolysis and that in its absence, infected mice are more resistant to fat loss. Lastly, we identified two host-protective functions for adipocyte lipolysis: prolonged host survival and transient reduction of AT parasite burden.

## Results

### *T. brucei* infection induces loss of AT weight and adipocyte area

Natural and experimental infections with *T. brucei* parasites induce weight loss, which includes a significant loss of fat mass. To study the dynamics of this process, C57BL/6J mice infected with pleomorphic *T. brucei* and, at different time-points post-infection, animals were sacrificed and the gonadal AT depot (i.e. the largest visceral AT depot) was harvested, weighed and processed for histology. Infected mice showed a progressive loss of gonadal AT weight. By day 9 post-infection a 30% loss of gonadal AT mass was observed, which progressed to a 54% and 65% loss by days 16 and 30 post-infection, respectively **(Figure 1A)**. To assess whether the observed loss of AT weight was associated with a reduction in adipocyte size, we analyzed Hematoxylin & Eosin - stained sections of the gonadal AT at different time-points post-infection. A progressive reduction in lipid droplet size was observed concomitant with a previously described accumulation of inflammatory infiltrates^6^ **(Figures 1B and 1C)**. Quantification of lipid droplet area revealed an average reduction of 16%, 37%, 61%, and 69%, by days 6, 10, 16 and 30 post-infection, respectively **(Figure 1C)**.

**Figure 1.**
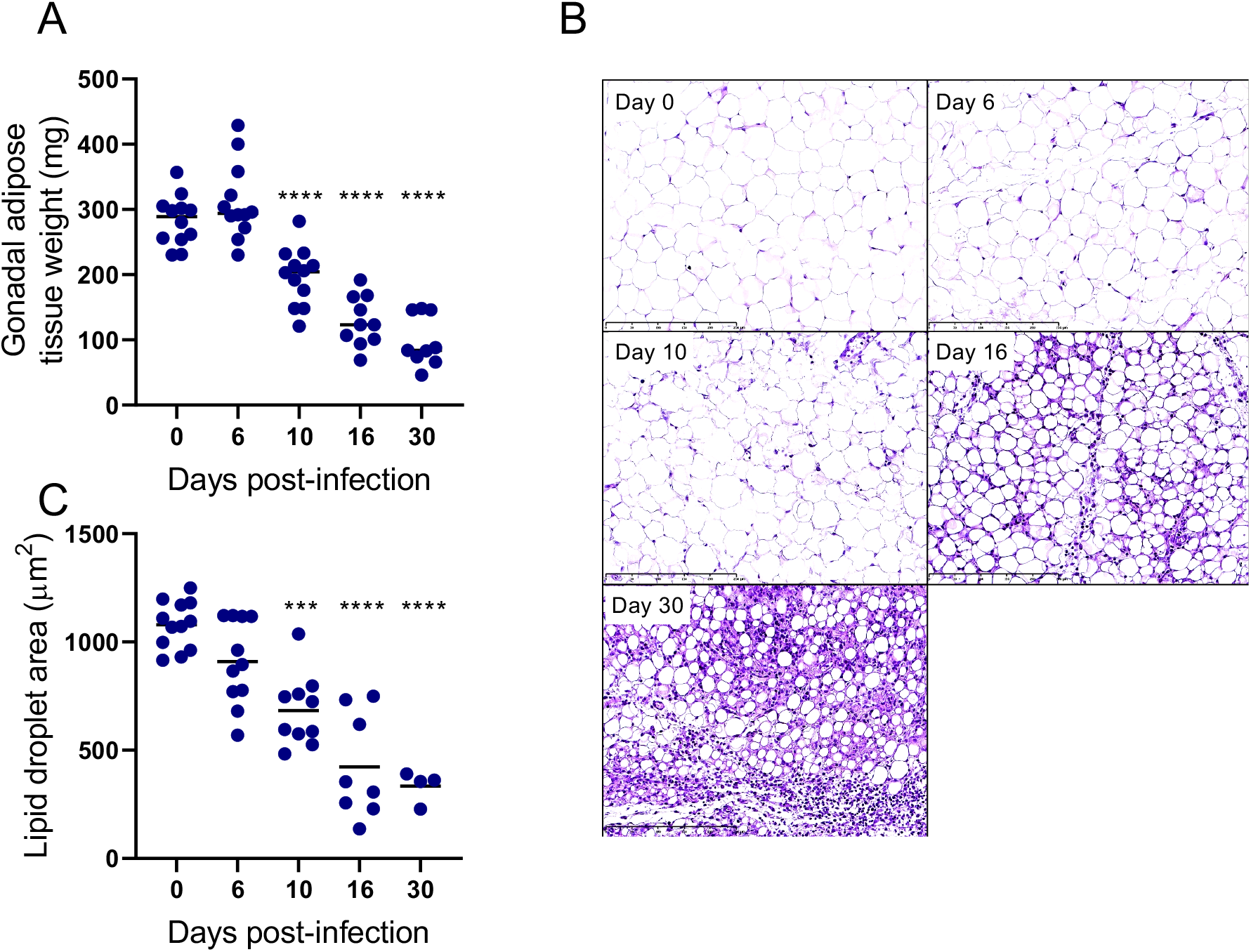
Dynamics of adipose tissue weight and adipocyte size during infection. **(A)** Weight of gonadal adipose tissue throughout infection. **(B)** Representative hematoxylin & eosin micrographs of gonadal AT at different time-points post-infection. Scale bar length = 250 μm, magnification = 20x. **(C)** Quantification of lipid droplet area of the gonadal AT at different time-points post-infection. (n = 4-12 mice per group, pooled data from 2 independent experiments). Statistical analysis was performed with one-way ANOVA using Sidak’s test for multiple comparisons. *, P<0.05; **, P<0.01; ***, P<0.001; ****, P<0.0001.

Overall, these data show that during infection white AT undergoes a progressive weight loss that is correlated with a reduction of individual adipocyte sizes. In turn, This imbalance appears to favor the mobilization of stored intracellular fat over lipogenesis, leading to a reduction in adipocyte volume.

### Adipocyte lipolysis is increased during a *T. brucei* infection

Next, we sought to identify whether loss of fat mass during *T. brucei* infection was a result of increased adipocyte lipolysis. Gonadal AT from infected and control mice was harvested at different time-points post-infection and incubated *ex vivo* to allow for the release of lipolytic products (i.e. fatty acids and glycerol), which were subsequently quantified and normalized to the total amount of protein in the tissue. A significant increase in free/non-esterified fatty acid (NEFA) release was observed from the gonadal AT of mice infected for 6 and 9 days (**Figure 2A)**. Conversely, by day 16 post-infection, a decrease in the release of fatty acids from the AT these mice relative to non-infected controls was observed **(Figure 2A)**. This later reduction in adipocyte lipolysis stems partially from a normalization bias that occurs at later time points of infection, as the AT becomes comparatively more protein rich as lipid stores are depleted and immune infiltration progresses **(Supplementary Figure 1)**.

**Figure 2.**
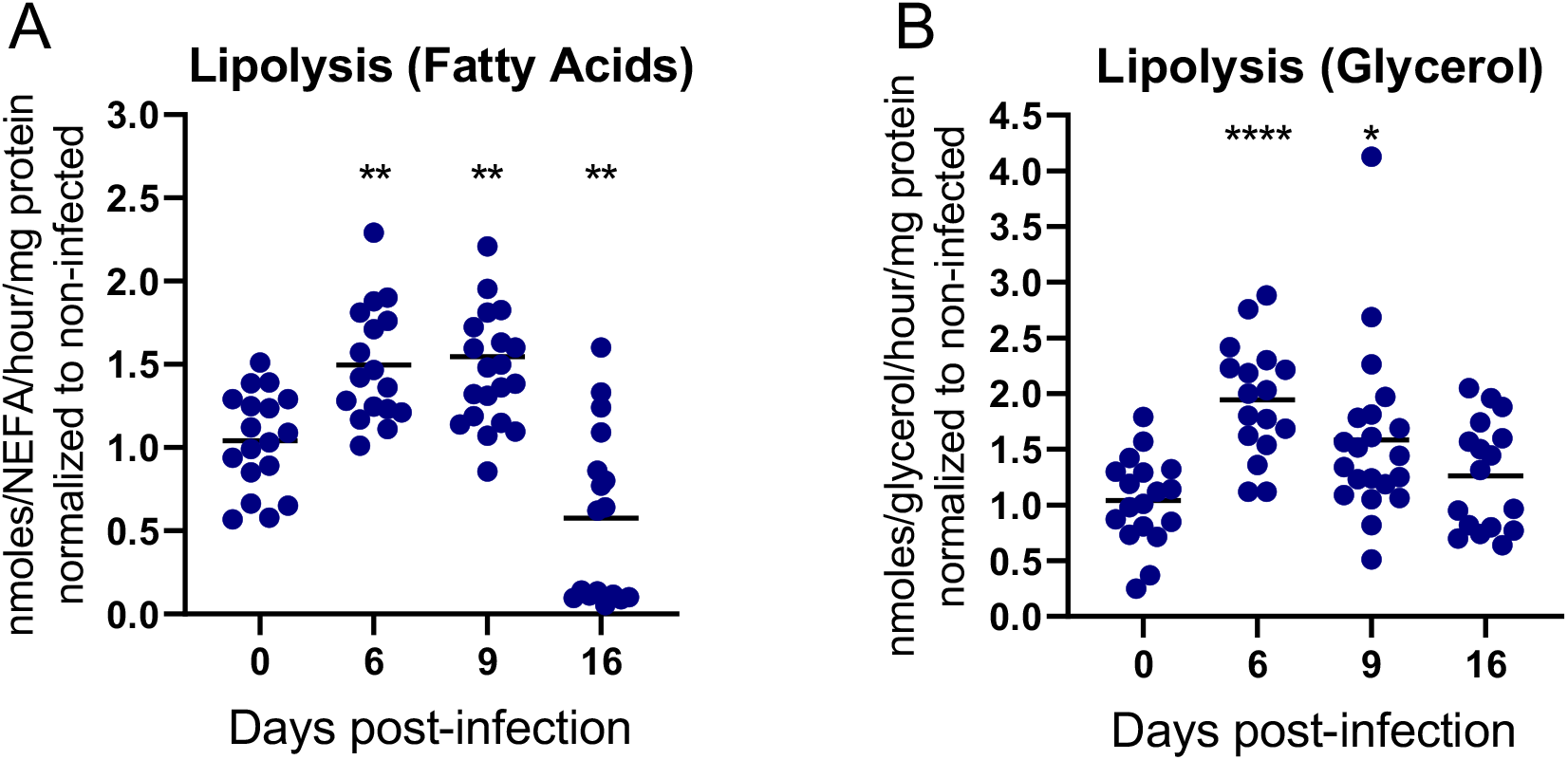
Increased lipolysis during a *T. brucei* infection. Release of **(A)** fatty acids and **(B)** glycerol from AT explants. (n = 17-18 mice per group, pooled data from 4 independent experiments). Statistical analysis was performed with one-way ANOVA using Sidak’s test for multiple comparisons. *, P<0.05; **, P<0.01; ***, P<0.001; ****, P<0.0001.

Likewise, an increase in glycerol release was observed by days 6 and 9 post-infection, but not by day 16 post-infection **(Figure 2B)**. On average the fold increase was for glycerol was higher than that observed for fatty acids. This difference may in part be due to glycerol secreted by the parasite, as significant amounts of glycerol are produced by *T. brucei* during glycolysis, especially under hypoxic conditions^17,18^. In line with this hypothesis, *T. brucei* cultures show a reduction of fatty acid concentration (i.e. C8 and longer) while accumulating glycerol in a dose dependent manner **(Supplementary Figure 2)**.

These results show that adipocyte lipolysis is activated during a *T. brucei* infection.

### Infection-induced lipolysis requires immune cues

Next we sought to identify the upstream signals responsible for activation of adipocyte lipolysis. In other contexts, adipocyte lipolysis can be induced by a myriad of distinct stimuli, including immune, neuronal and hormonal signals^7^.

First, we tested whether adipocyte lipolysis activation was mediated by modulation of sympathetic tone. To test whether sympathetic innervation contributed to *T. brucei* induced fat loss, we infected mice chemically sympathectomized with 6-hydroxydopamine (6-OHDA), a neurotoxic analogue of dopamine. This compound acts by damaging nerve terminals of sympathetic neurons, preventing norepinephrine release^19^. No significant differences were observed in fat mass loss between 6-OHDA treated and sham treated infected mice **(Supplementary Figure 3A)**, suggesting that changes in sympathetic tone are not involved in loss of fat mass during a *T. brucei* infection.

Second, we tested whether infected animals were losing fat mass because of their feeding behavior. Indeed, we have previously described that there is a transient disease-induced hypophagia observed around the first peak of parasitemia (i.e. days 4 to 10 post-infection)^5^. To test if hypophagia is sufficient to induce adipocyte lipolysis, we employed two distinct experimental setups of controlled food intake: fasting & re-feeding setup and a paired-feeding setup. In the fasting and re-feeding setup, we synchronized the feeding behavior of infected and non-infected groups by fasting them for 7 hours during peak activity hours, followed by a 2 hours re-feeding period prior to being euthanized **(Figure 3A)**. In these conditions, infected mice continued to show the same dynamic increase in the rate of adipocyte lipolysis during first peak of parasitemia, with a reduction after day 10-12 post-infection (**Figures 3B-C)**. In the paired feeding setup **(Figure 3D)**, infected mouse are allowed to feed *ad libitum* (i.e. unrestricted access to food) and the amount of consumed food of each mouse was weighed. These food intakes determined the amount of food provided to each pair-fed non-infected control the following day **(Figure 3D-E)**. In this setup, infected mice showed again higher adipocyte lipolysis than their respective non-infected controls. Moreover, the non-infected paired-fed control did not show a significant increase in adipocyte lipolysis **(Figures 3F-G)**, suggesting that disease induced-hypophagia is not sufficient to induce adipocyte-lipolysis during a *T. brucei* infection.

**Figure 3.**
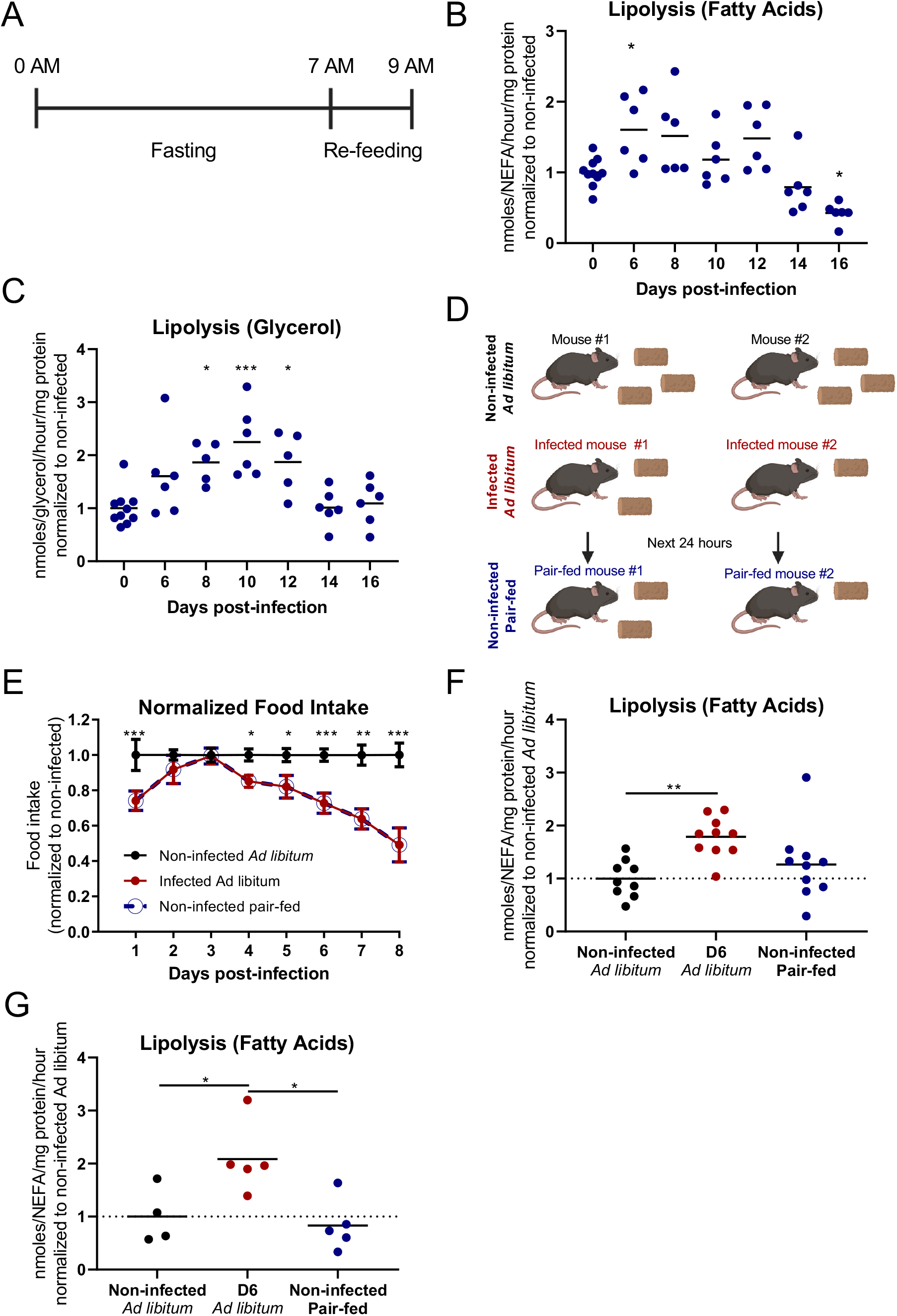
Changes in feeding behavior are not responsible for *T. brucei* induced adipocyte lipolysis. **(A)** Schematic representation of the fasting and re-feeding experimental layout. Release of **(B)** fatty acids and **(C)** glycerol from gonadal AT explants of infected and non-infected WT mice, normalized to non-infected mice. **(D)** Schematic representation of the paired-feeding experimental layout. **(E)** Daily food intake normalized to non-infected *ad libitum* fed mice. **(F-G)** Release of fatty acids from gonadal AT explants of infected and non-infected WT mice, normalized to ad libitum fed non-infected mice. **(B-C)** n= 5-10 mice per group. **(E-G)** n=4-9 mice per group. Statistical analysis was performed with **(B-C and F-G)** One-way and **(E)** Two-way ANOVA using Sidak’s test for multiple comparisons. *, P<0.05; **, P<0.01; ***, P<0.001; ****, P<0.0001. **(E-F)** Pooled data from two independent experiments.

As *T. brucei* induces both a strong systemic and local (i.e. within the AT) immune response, next we questioned whether immune activation was required for lipolysis upregulation. To test this hypothesis, we infected recombination activation gene-2 deficient (*Rag2*^-/-^) mice which lack both B and T cells. These immunosuppressed mice present hyperparasitemia **(Figure 4A)** and usually succumb to infection by day 15 post-infection^20^.

**Figure 4.**
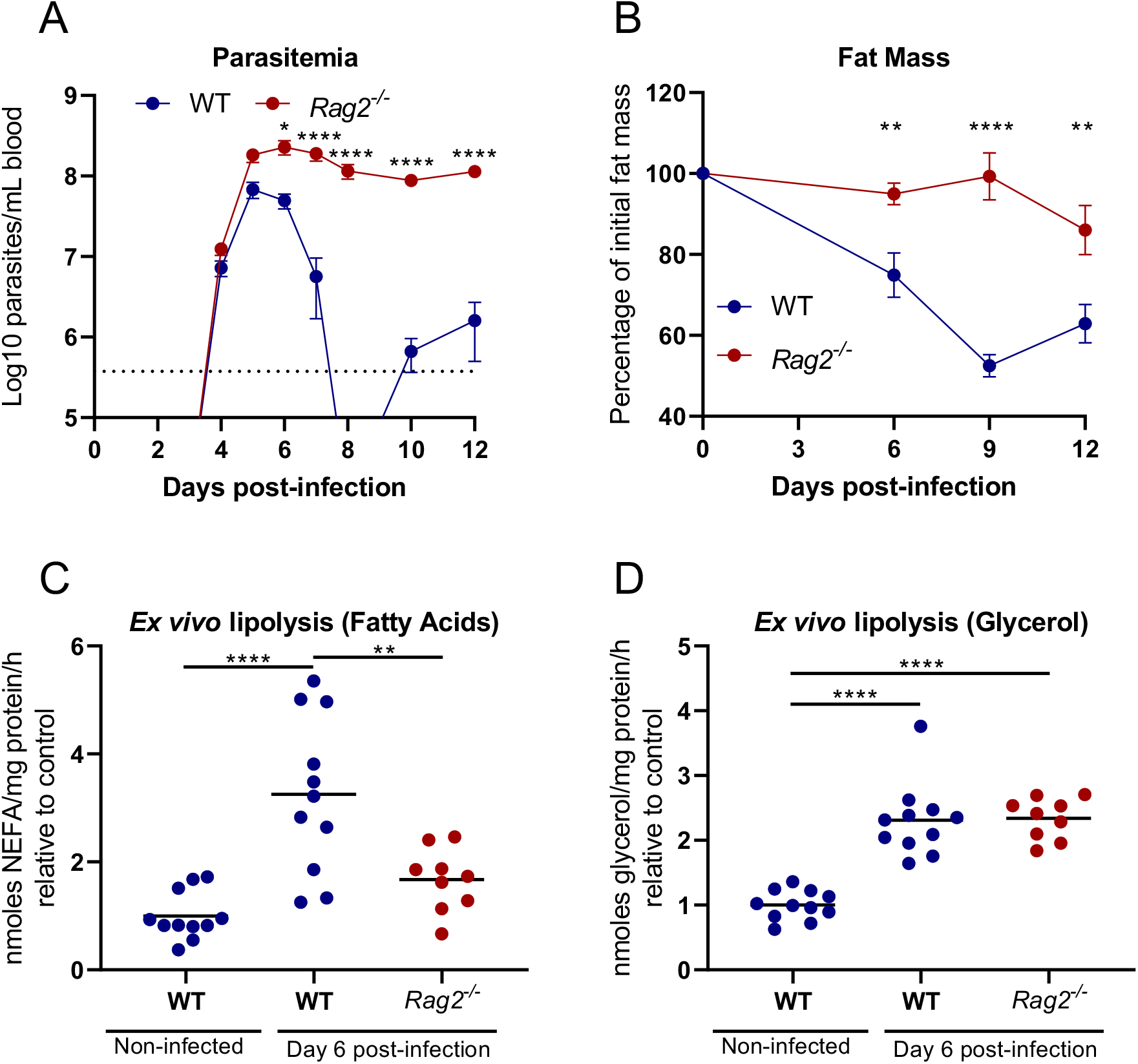
*T. brucei* induced loss of fat mass and adipocyte lipolysis is dependent on an adaptive immune response. **(A)** Parasitemia of infected WT and Rag2^-/-^ mice. **(B)** Fat mass relative to baseline of infected *Rag2*^*-/-*^ and WT controls. Release of **(C)** fatty acids and **(D)** glycerol from gonadal AT explants of infected WT and *Rag2*^*-/-*^ mice normalized to non-infected WT controls. Error bars represent the SEM. (**(A-B)** n= 6-7 and **(C-D)** n= 9-11 mice per group, pooled data from 2 independent experiments). Statistical analysis was performed with **(A-B)** Two-way ANOVA and (**C-D)** One-way ANOVA using Sidak’s test for multiple comparisons. *, P<0.05; **, P<0.01; ***, P<0.001; ****, P<0.0001.

Longitudinal body composition analysis of infected *Rag2*^*-/-*^ mice revealed a clear reduction in the loss of overall fat mass when compared to infected WT controls **(Figure 4B)**. Indeed, by day 6 and 9 post-infection while WT mice presented a 25% and 48% reduction in total fat mass, *Rag2*^*-/-*^ mice showed no significant loss of fat mass. As infection progressed, *Rag2*^*-/-*^ mice showed a modest loss of fat mass. Specifically, by day 12 post-infection *Rag2*^*-/-*^ mice had lost 14% of total fat mass, significantly less than the 37% loss of fat mass exhibited by infected WT mice at the same time.

To assess whether this higher retention of fat mass in infected *Rag2*^*-/-*^ mice correlated with a failure to upregulate adipocyte lipolysis, we quantified the rate of release of fatty acids from gonadal AT explants of infected *Rag2*^*-/-*^ mice and compared it to both infected and non-infected WT controls **(Figures 4C-D)**. While the rate of fatty acid release in infected WT mice increased by around 3.25-fold, relative to non-infected mice, the rate of fatty acid release in infected *Rag2*^*-/-*^ mice increased by only 1.67-fold **(Figure 4C)**. While AT of infected *Rag2*^*-/-*^ showed a similar release of glycerol to AT of infected WT mice **(Figure 4D)**, this probably stems from an increase in parasite-secreted glycerol due to increased AT parasite burdens^6^. Interestingly, mice deficient for TNF-α, a cytokine classically associated with *T. brucei*-induced wasting and a known activator of adipocyte lipolysis showed only a small delay in fat mass retention **(Supplementary Figure 3B)**, suggesting that additional inflammatory factors are involved in fat mass loss and lipolysis induction.

Together, these data show that activation of adipocyte lipolysis is independent of mouse feeding behavior and it is not mediated by sympathetic nervous system nor by TNFα alone. Instead, we observed that loss of fat mass during a *T. brucei* infection is strongly dependent on the presence of T and/or B lymphocytes,

### Loss of adipocyte volume is dependent on adipocyte ATGL activity

After establishing that adipocyte lipolysis is increased during a *T. brucei* infection, we questioned whether we could revert the fat loss phenotype by genetically preventing lipolysis from being activated. For this, we used adipocyte-specific ATGL deficient mice (*Adipoq*^*Cre/+*^*-Atgl*^*fl/fl*^), which express a Cre recombinase under the control of the adiponectin promoter and have two loxP sites flanking the *Pnpla2* (ATGL) gene. Thus, these mice have no expression of ATGL in cells that express adiponectin (i.e. adipocytes)^21,22^.

Using this model, we first asked if lipolysis could still be activated during an infection by *T. brucei*. Using the same lipolytic assay described above, gonadal AT was collected and levels of released fatty acids and glycerol were measured *ex vivo*.

Infected *Adipoq*^*Cre/+*^*-Atgl*^*fl/fl*^ (KO) mice showed a significantly lower secretion of fatty acids and glycerol than infected *Atgl*^fl/fl^ (WT) littermate mice **(Figures 5A-B)**, indicating that activation of ATGL is essential for lipolysis induced during a *T. brucei* infection.

**Figure 5.**
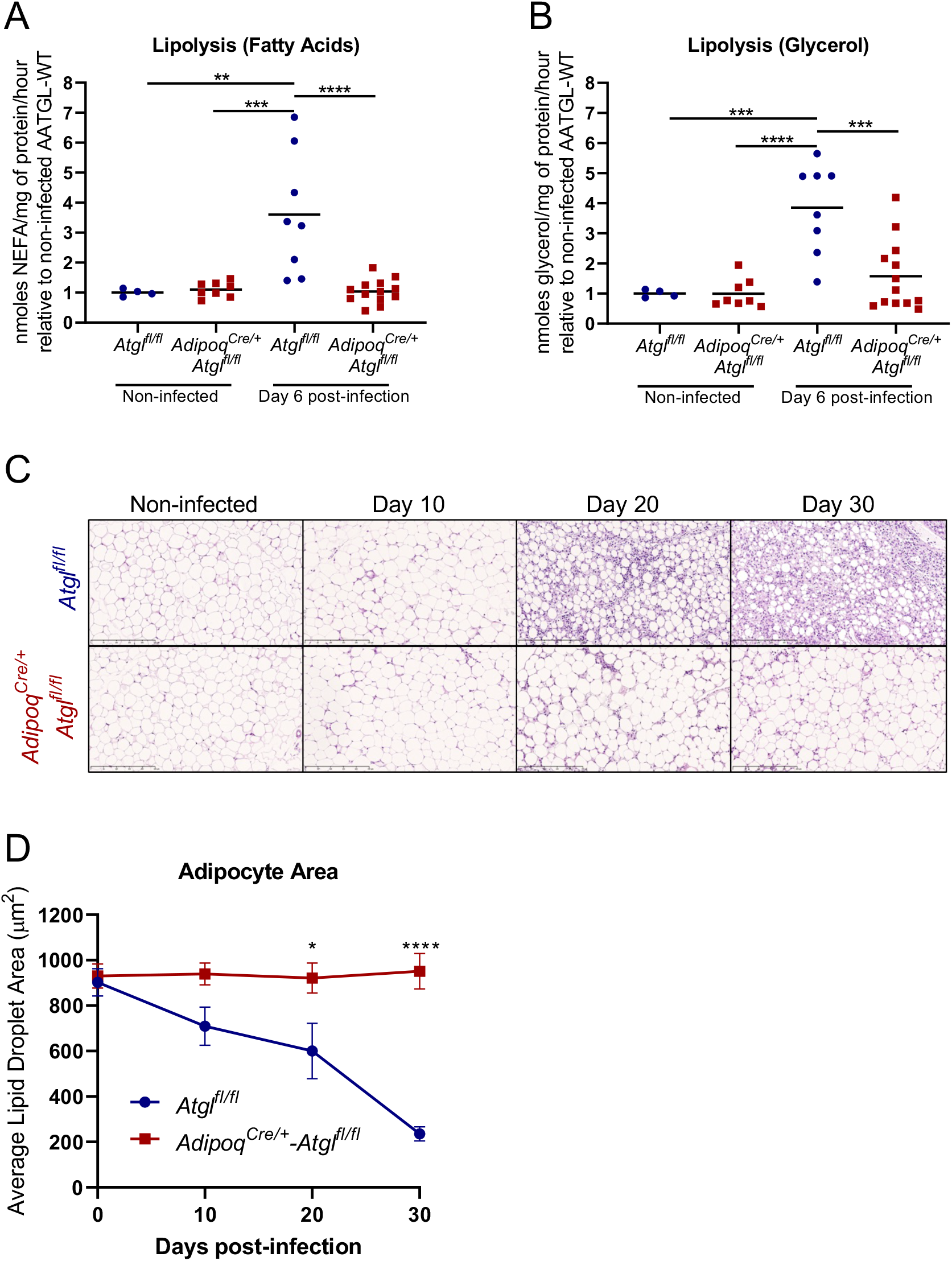
ATGL deficiency in adipocytes prevents *T. brucei* induced lipolysis and loss of adipocyte area. Release of **(A)** fatty acids and **(B)** glycerol from gonadal AT explants of infected and non-infected *Atgl*^fl/fl^ (WT) and *Adipoq*^*Cre/+*^*-Atgl*^fl/fl^ (KO) mice. **(C)** Representative gonadal AT hematoxylin and eosin micrographs (20x magnification, scale bar 250 μm). **(D)** Quantification of lipid droplet area of the gonadal AT at different time-points post-infection. Error bars represent the SEM (**(A-B)** n= 4-12 and **(D)** n=3-6 mice per group). Statistical analysis was performed with **(A-B)** One-way ANOVA or **(D)** Two-way ANOVA using Sidak’s test for multiple comparisons. *, P<0.05; **, P<0.01; ***, P<0.001; ****, P<0.0001. **(A-B)** pooled data from two independent experiments.

Consistently, histology analysis of sections of gonadal adipose tissue revealed that *Adipoq*^*Cre/+*^*-Atgl*^*fl/fl*^ mice maintain a stable average adipocyte size (*i*.*e*. 900 μm^2^) while *Atgl*^fl/fl^ mice experience a progressive reduction in adipocyte size of up to 75% by day 30 post-infection, indicating that ATGL-dependent lipolysis is central in *T. brucei* induced adipocyte size reduction **(Figures 5C-D)**.

Overall, these data show that ATGL-mediated lipolysis in adipocytes is the main mechanism of fat mass loss during *T. brucei* infection.

### Loss of adipocyte ATGL activity impacts body composition

Next, we questioned how ATGL-deficiency may impact mouse whole body composition. Specifically, we hypothesized that in the absence of ATGL activity infected mice would be unable to mobilize energy stored in the form of TAG within adipocytes. In turn, this could potentially be compensated by alternative catabolic pathways such as skeletal muscle proteolysis, which would precipitate the hallmark wasting phenotype observed during natural *T. brucei* infections. To assess this, we performed a longitudinal study of body composition by nuclear magnetic resonance, which allowed us to follow total body weight, lean mass, free fluid and fat mass.

Total body weight of *Atgl*^fl/fl^ and *Adipoq*^*Cre/+*^*-Atgl*^*fl/fl*^ followed a similar profile, with an initial loss of approximately 5% during the first peak of parasitemia (i.e. days 5-8 post-infection), followed by a rapid recovery and relative stability until day 20 post-infection **(Figure 6A)**. Afterwards, total body weight declined for both WT and KO mice, with a trend for *Adipoq*^*Cre/+*^*-Atgl*^*fl/fl*^ to show lower total weight during the latest stages of infection (i.e. by day 30 post-infection) **(Figure 6A)**.

**Figure 6.**
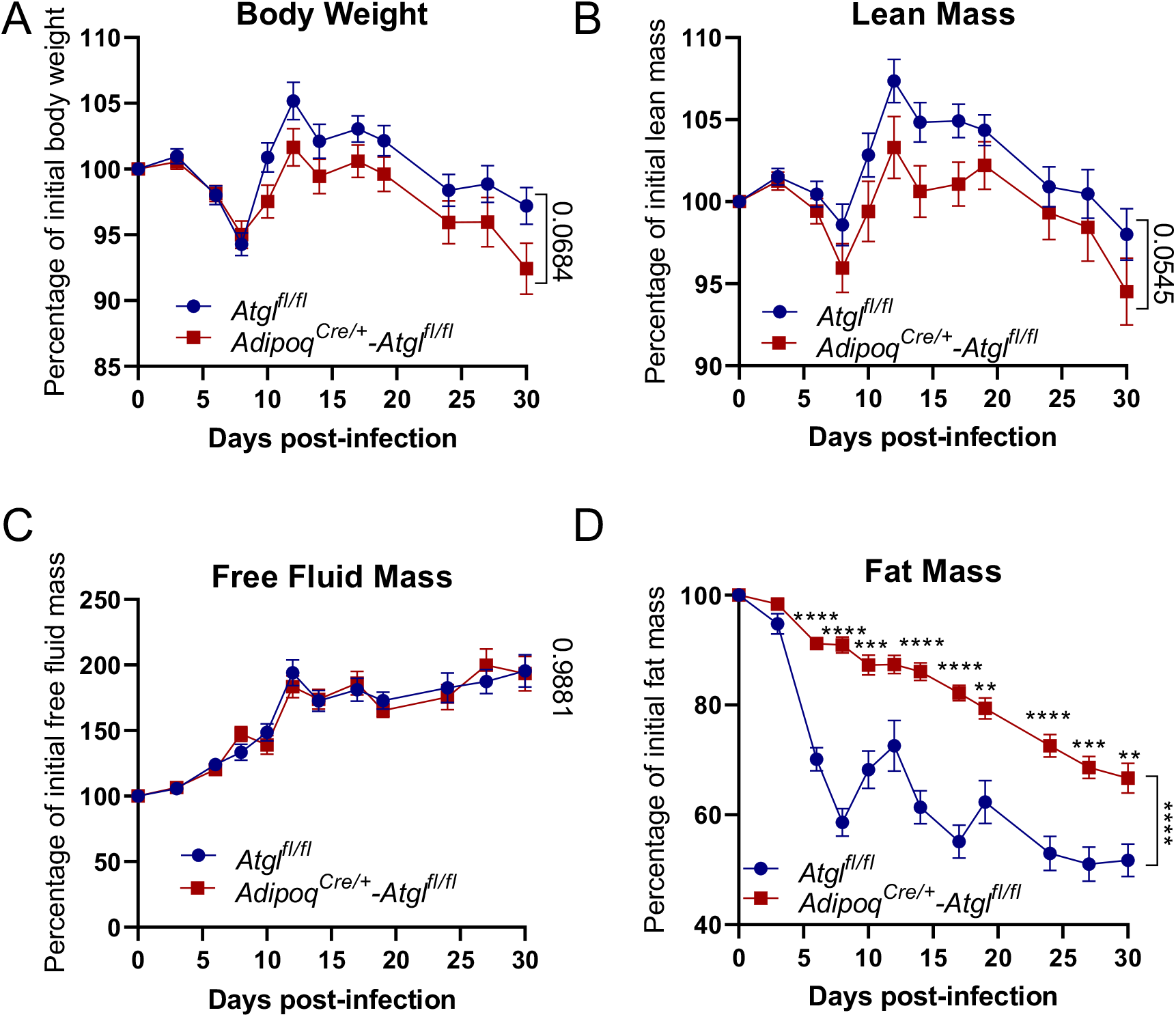
Effect of adipocyte-specific ATGL deficiency in body composition and skeletal muscle pathology. **(A)** Total body weight, **(B)** lean mass, **(C)** free fluid mass and **(D)** fat mass relative to baseline. Error bars represent the SEM (**(A-D)** n= 27-32 mice per group). Statistical analysis was performed with a mixed-effects Two-way ANOVA using Sidak’s test for multiple comparisons. *, P<0.05; **, P<0.01; ***, P<0.001; ****, P<0.0001. **(A-D)** pooled data from four independent experiments.

Similar to total body weight, a trend for comparatively reduced lean mass retention in *Adipoq*^*Cre/+*^*-Atgl*^fl/fl^ mice was observed **(Figure 6B)**. This trend was observed as early as day 8 post-infection, and was kept throughout infection in spite of the overall increase in total lean mass observed in the following days. This increase in lean mass comparative to baseline peaked by day 12 post-infection and likely stems from hepatomegaly and splenomegaly observed during *T. brucei* infections^23^.

During infection, a clear increase in free fluid mass was observed in both WT and KO mice **(Figure 6C)**, which was likely due to the formation of widespread edema that stems from the increased vascular leakage known to occur during *T. brucei* infection^24^. Importantly, significant increases of free fluid masses may act as a confounding factor when using total weight as a parameter for evaluating body condition of *T. brucei* infected mice.

Lastly, and as expected, *Adipoq*^*Cre/+*^-*Atgl*^fl/fl^ mice presented significantly increased retention of fat when compared to *Atgl*^fl/fl^ mice **(Figure 6D)**. By day 8 post-infection, while *Atgl*^fl/fl^ mice lost over 40% of total fat mass, *Adipoq*^*Cre/+*^*-Atgl*^fl/fl^ mice presented only a modest 8% loss **(Figure 6D)**. Afterwards, *Adipoq*^*Cre/+*^*-Atgl*^fl/fl^ mice steadily lost fat mass, reaching an average 33% loss by day 30 post-infection – significantly less than the 49% experienced by *Atgl*^fl/fl^ mice. Moreover, *Adipoq*^*Cre/+*^*-Atgl*^fl/fl^ mice showed high AT mass retention until the latest stages of infection, as necropsy of moribund *Adipoq*^*Cre/+*^*-Atgl*^fl/fl^ mice revealed significantly higher weight of gonadal AT depots than those of *Atgl*^fl/fl^ mice **(Supplementary Figure 4)**. These data suggest that both ATGL-dependent and ATGL-independent mechanisms contribute to fat loss during a *T. brucei* infection.

Overall, these data suggest that ATGL activity in adipocytes during infection promotes loss of fat mass and potentially exerts a mild protective role in lean mass retention.

### Adipocyte lipolysis prolongs host survival and reduces AT parasite burden

Having established that adipocyte ATGL plays a central role in fat mass loss during a *T. brucei* infection, next we asked how activation of lipolysis affected disease progression. We hypothesized that given that *Adipoq*^*Cre/+*^*-Atgl*^fl/fl^ infected mice do not undergo loss of fat mass, animals would be more resistant to infection and would survive longer. Surpisingly, *Adipoq*^*Cre/+*^*-Atgl*^*fl*/fl^ mice succumbed earlier to infection than their WT littermate controls **(Figure 7A)**, suggesting a protective role for ATGL-mediated lipolysis during infection. This increased susceptibility of *Adipoq*^*Cre/+*^*-Atgl*^fl/fl^ mice was not due to an increase in parasitemia, as absence of this lipase had no significant effect on the control of the first peak of parasitemia or during the latest stages of infection (i.e. 10 days preceding death) **(Figure 7B)**.

**Figure 7.**
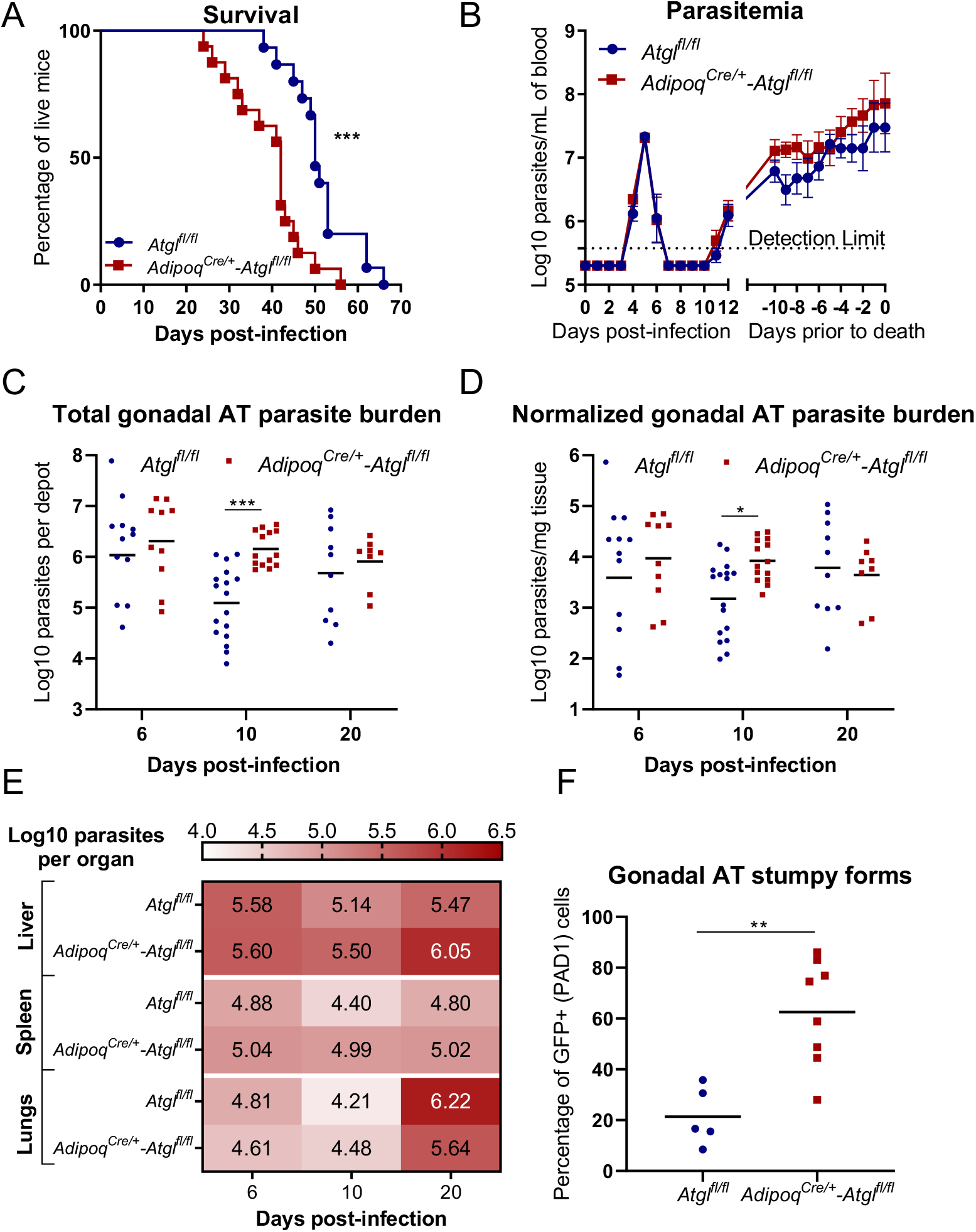
Effect of adipocyte-specific ATGL deficiency in infection. **(A)** Survival curves and **(B)** Log10 parasitemia of *T. brucei* infected *Atgl*^fl/fl^ and *Adipoq*^*Cre/+*^-*Atgl*^fl/fl^ mice. Log10 number of *T. brucei* parasites at different time-points post-infection per **(C)** depot or **(D)** milligram of gonadal AT of *Atgl*^fl/fl^ and *Adipoq*^*Cre/+*^-*Atgl*^fl/fl^ mice. **(E)** Heatmap of number of *T. brucei* parasites in the liver, spleen and lungs of *Atgl*^fl/fl^ and *Adipoq*^*Cre/+*^*-Atgl*^fl/fl^ mice at different time-points post-infection. **(F)** Percentage of PAD1+ parasites extracted from the gonadal AT of *Atgl*^fl/fl^ and *Adipoq*^*Cre/+*^*-Atgl*^fl/fl^ mice at day 6 post-infection. Statistical analysis was performed with **(A)** Log-rank (Mantel-Cox) test, **(B-E)** two-way ANOVA using Sidak’s test for multiple comparisons and **(F)** unpaired t test. *, P<0.05; **, P<0.01; ***, P<0.001; ****, P<0.0001. **(A-B)** n= 15-16 mice per group, pooled data from 2 independent experiments. **(C-D)** n= 8-17 mice per group, pooled from 2 independent experiments. **(E)** n=4-7 mice per group from a single experiment. **(F)** n=5-8 mice per group from a single experiment.

To understand how adipocyte lipolysis contributes to prolonged host survival, we questioned whether adipocyte lipolysis *per se* is able to modulate the number of parasites. As previously shown, *Adipoq*^*Cre/+*^*-Atgl*^fl/fl^ mice did not show different blood parasite burdens when compared to WT littermate controls. This indicates that adipocyte lipolysis is insufficient to modulate number of parasites on a systemic level, but it does not exclude the existence of tissue-specific effects. Quantification of parasite load in the gonadal AT (by qPCR) revealed that the parasite number is 10-fold higher in *Adipoq*^*Cre/+*^*-Atgl*^fl/fl^ mice that WT littermate controls at day 10 post-infection **(Figure 7C)**. By day 20 post-infection, this difference in parasite load was no longer detectable, which correlates with the fact that lipolysis is no longer activated at stage of infection **(Figure 2)**. To remove any bias due to a higher AT mass typically observed in *Adipoq*^*Cre/+*^*-Atgl*^fl/fl^ compared to *Atgl*^fl/fl^ mice (i.e. due to loss of AT mass in *Atgl*^fl/fl^ mice, we normalized the number of parasites to total tissue mass **(Figure 7D)**. After this normalization, *Adipoq*^*Cre/+*^*-Atgl*^fl/fl^ mice present on average a significant 7-fold increase in the number of parasites in the gonadal AT compared to *Atgl*^fl/fl^ controls. This higher parasite load appears to be restricted to the AT, as no significant differences in parasite load were found between the livers, spleens of lungs of *Atgl*^fl/fl^ and *Adipoq*^*Cre/+*^*-Atgl*^fl/fl^ mice **(Figure 7E)**.

During a *T. brucei* infection, the total parasite load depends in part on a quorum sensing mechanism that triggers differentiation of replicative forms into non-replicative stumpy forms^25^. We hypothesized that the higher parasite load detected in gonadal AT of *Adipoq*^*Cre/+*^*-Atgl*^fl/fl^ mice could be due to reduced parasite differentiation into the non-replicative stumpy form. To quantify the proportion of replicative and non-replicative forms in AT population, we infected *Atgl*^fl/fl^ and *Adipoq*^*Cre/+*^*-Atgl*^fl/fl^ mice with a parasite reporter cell line that expresses GFP when differentiation is triggered (PAD1::GFP reporter). We isolated parasites from the gonadal AT at day 6 post-infection (i.e. peak of AT parasite burden) and quantified the frequency of GFP positive cells by flow cytometry. Contrary to our hypothesis, parasites extracted from the AT of *Adipoq*^*Cre/+*^*-Atgl*^fl/fl^ mice presented a higher percentage of stumpy forms (i.e. PAD1::GFP expressing parasites) **(Figure 7F)**. While the population of non-replicative stumpy forms in *Atgl*^fl/fl^ mice was approximately 20%, in *Adipoq*^*Cre/+*^*-Atgl*^fl/fl^ mice was 60%, indicating that parasite differentiation is not inhibited in the absence of adipocyte lipolysis.

Together, these data suggest that adipocyte lipolysis induces a local host-protective effect, limiting the number of parasites. It is possible that released fatty acids and glycerol exert an immunomodulatory effect that increases the trypanocydal capacity of the local immune response. Alternatively, these lipolytic products may directly modulate the parasite’s cell cycle or exert a direct trypanocydal effect (e.g. lipotoxicity).

## Discussion

In this work, we sought to gain mechanistic insight on how a *T. brucei* infection leads to loss of fat mass, and in turn how the pathways governing this loss of fat mass may affect disease progression and outcome. During the course of this study, we found that loss of fat mass and adipocyte size requires activation of neutral lipolysis up to day 9 post-infection, particularly through the activity of the neutral lipolysis rate-limiting enzyme ATGL. Moreover, we found evidence that this activation of lipolysis is tightly dependent on immune activation, as *Rag2-/-* mice despite having a higher parasite burden, show significantly lower activation of lipolysis and loss of fat mass than WT controls. Importantly, we showed that ATGL-dependent adipocyte lipolysis has an overall host-protective role in the context of a *T. brucei* infection, as in its absence, mice succumb earlier to infection and show higher AT parasite burden during the period of infection where lipolysis would be activated.

### Impact of ATGL-dependent adipocyte lipolysis on body fat

In this work we have shown that during a *T. brucei* infection adipocytes become progressively smaller, in a process that involves an increase in ATGL-dependent lipolysis. Other studies have shown a similar role for ATGL in the loss of fat mass. Specifically, full ATGL-deficient mice are protected from fat mass loss induced by Lewis lung carcinoma or B16 melanoma cells^14^. Moreover, burn injury-induced loss of body weight and adipocyte volume was partially prevented in *Adipoq*^*Cre/+*^-*Atgl*^fl/fl^ mice^26^. Interestingly, in a model of lymphocytic choriomeningitis virus infection, where significant loss of AT and total weight occur, *Adipoq*^*Cre/+*^-*Atgl*^fl/fl^ mice were not protected from loss of total weight^27^. This suggests the relative relevance of ATGL in fat mass retention in infectious disease is pathogen-specific.

Increased adipocyte lipolysis is likely one among metabolic alterations occurring simultaneously. Indeed, work in the early 1980s using rabbits experimentally infected with *T. brucei* established a link between hyperlipidemia and reduced lipoprotein lipase (LPL) activity due to increased TNF-α levels^28^. Here, inhibition of LPL prevented the export of lipids contained in lipoproteins from circulation and into the AT, leading to decreased lipogenesis.

Elevated TNF-α levels are often found in pathological conditions where wasting phenotypes develop^29,30^. Moreover, treatment of several cell types, including adipocytes, with this cytokine promotes the upregulation of catabolic processes in detriment of anabolic ones, a phenotype often termed as cellular cachexia^31^. Although never formally tested, this collective body of work led to the logic conclusion that inhibition of lipogenesis may be central in the progressive wasting phenotype observed in during *T. brucei* infections. Recently, a greater appreciation of the complex inflammatory profile leading to the development of cachexia has re-framed the role of TNF-α as one of many inflammatory cytokines in the development and progression of wasting syndromes^32^.

Although it remains to be elucidated exactly how relevant LPL activity is in the loss of fat mass during a *T. brucei* infection, our data suggests that TNF-α is not required to induce fat loss. Consistently, TNF-α deficient mice, which should retain significant LPL activity, showed only a small delay in loss of fat mass **(Supplementary Figure 3B)**. Indeed, our data show that fat loss and adipocyte volume reduction require the activation of adipocyte lipolysis **(Figure 5)**. This dominance of increased adipocyte lipolysis over reduced lipogenesis in fat loss has also been reported in the context of CAC^33^.

Ablation of adipocyte-ATGL significantly impaired loss of whole-organism fat mass, however it was unable to completely abolish it **(Figure 6D)**. On the one hand, even in absolute absence of adipocyte-lipolysis, a significant loss of total fat mass would still be expected, as most cells are capable of storing fats within cytosolic lipid droplets, which may then be consumed during infection. On the other hand, it is likely that even in the absence of adipocyte-ATGL during a *T. brucei* infection, TAG may still be hydrolyzed at a slower rate by HSL. This lipase is able to hydrolyze TAG, despite having 10-fold higher affinity for DAG^34^. Moreover, if at any point acid lipolysis is activated in adipocytes during infection, lysosomal acid lipase would be able to fully catalyze the conversion of TAG into free fatty acids and glycerol without requiring the activity of any neutral lipase^35^.

Altogether, our data show that activation of adipocyte lipolysis is a central, but not exclusive, pathway in fat loss and emaciation during *T. brucei* infections.

### Induction of adipocyte lipolysis

In this work, we attempted to isolate the specific signal(s) that induce adipocyte lipolysis during a *T. brucei* infection. Our data suggest that modulation of sympathetic tone or changes in feeding behavior are insufficient to drive *T. brucei*-induced adipocyte lipolysis **(Figure 3 and Supplementary Figure 3A)**. In contrast, a recent report found that changes in food intake in male mice correlate with weight loss^36^. This apparent inconsistency of results could be due to the fact that in this study the authors focused on total weight loss, whereas we characterized lipolysis induction and subsequent loss of fat loss in the AT.

On the other hand, T and B cell-secreted cytokines (or secreted by other cells after T and/or B cell-dependent activation) are better candidates as inducers of adipocyte lipolysis. This immune-centric hypothesis is supported by the reduced loss of fat mass observed in infected *Rag2*^*-/-*^ mice **(Figure 4B)** and by their low *ex vivo* AT lipolysis upon infection **(Figure 4C)**.

A multitude of immune factor, notably cytokines, have been shown to have a pro-lipolytic effect. These include interleukin (IL)1, IL6, IL15, IL17A, TNF-α, growth/differentiation factor (GDF)15, leukemia inhibitory factor (LIF) as well as type I and type II IFNs^32,37^. A recent study found that IL-17 drives weight loss in male mice^36^. In a complex inflammatory scenario, such as a *T. brucei* infection, it is likely that multiple of these factors are simultaneously elevated. Accordingly, it is feasible that the observed upregulation of lipolysis stems from a combined/synergistic effect of these cytokines rather than a dominant effect of any of these candidates. The precise nature of the pro-lipolytic factor(s) driving adipocyte lipolysis during a *T. brucei* infection remains to be elucidated.

### Effect of adipocyte lipolysis on pathology

Mice deficient for ATGL-dependent adipocyte lipolysis showed a significant reduction in survival when compared to WT littermate controls **(Figure 7A)**. These differences were not due to changes in parasitemia, as *Adipoq*^*Cre/+*^*-Atgl*^fl/fl^ mice were equally able as *Atgl*^fl/fl^ to clear the successive parasitemia peaks, suggesting that adipocyte lipolysis does not impact systemic parasite elimination **(Figure 7B)**.

Interestingly, albeit non-significant, a clear trend for *Adipoq*^*Cre/+*^*-Atgl*^fl/fl^ mice to present lower total body weight and importantly lean mass than *Atgl*^fl/fl^ was observed. These data suggest that in the absence of efficient adipocyte lipolysis mice undergo a more severe wasting phenotype. In turn, it is tempting to speculate that this more pronounced loss of body condition is sufficient to decrease *Adipoq*^*Cre/+*^-*Atgl*^fl/fl^ mice tolerance to disease, resulting in premature death.

### Local parasite burden control by adipocyte lipolysis

The abundant availability of fatty acids and glycerol in the AT milieu may provide the parasite with additional substrates for energy production through β-oxidation and glycolysis. It is thus puzzling that increasing the availability of these metabolites through upregulation of adipocyte lipolysis leads to a significantly reduced local parasite burden.

Our data show that when lipolysis is blocked, the number of parasites in adipose tissue is 10-fold higher than when lipolysis is active. We found that parasite differentiation from the replicative to the non-replicative stumpy forms is not impaired in adipose tissue when lipolysis is blocked, suggesting lipolysis may be limiting parasite growth through an alternative mechanism. An overall limitation of parasite population results in lower parasite densities, which ultimately prevents activation of the density-dependent parasite quorum sensing mechanism that governs non-replicative stumpy differentiation^38^.

A possible mechanism through which adipocyte lipolysis reduces parasite density relies on fatty acid induced lipotoxicity. In this scenario, when lipolysis is activated, more free fatty acids are released to the extracellular space, which are endocytosed by *T. brucei* and could be toxic if they exceed the parasite’s capacity to catabolize these species through β-oxidation^5^ or to anabolize them into relatively stable acylglycerol species stored into lipid droplets. This form of lipid-mediated parasite killing has been elegantly shown in *Toxoplasma gondii*, where excessive supplementation of unsaturated C18 and C16 fatty acids or inhibition of *T. gondii* diacylglycerol acyltransferase (DGAT)-1 (*i*.*e*. inhibiting lipid droplet formation) led to a marked reduction of doubling time and parasite viability^39^.

Although lipid droplet formation has is yet to be thoroughly addressed in mammalian stages of *T. brucei*, there is compelling evidence that their formation is essential for procyclic from survival^40^. Here, ablation of TbLpn, (Tb927.7.5450) a lipin that converts phosphatidic acid into diacylglycerol, leads to defective lipid droplet formation, mitochondrial abnormalities and reduced parasite viability under culture conditions^40^. Considering the impact of lipid droplet formation in both *T. brucei* procyclic forms and in mammalian infective *T. gondii*, it is tempting to speculate that similar lipotoxicity phenotypes may occur with mammalian stages of *T. brucei* in the interstitial spaces of the AT upon upregulation of adipocyte lipolysis.

Overall, further study on how adipocyte lipolysis improves host control over *T. brucei* may lead to the identification of new mechanisms that may be useful in our understanding not only of the pathology of trypanosomiasis, but whether this is a general mechanism of host defense against infectious diseases. Ultimately, identification of this disease mechanism can lead to the development of novel therapies against Human or Animal African trypanosomiasis.

## Methods

### Ethics statement

All experimental animal work was performed in accordance with the Federation of European Laboratory Animal Science Associations (FELASA) guidelines and was approved by the Animal Care and Ethical Committee of the Instituto de Medicina Molecular João Lobo Antunes (under the licenses AWB_2016_07_LF_Tropism and AWB_2021_11_LF_TrypColonization).

### Animals

C57BL/6J mice were purchased from Charles River Laboratories International. Genetically modified mice include: *Rag2*^*-/-*^ (IMSR_JAX:008449), *Tnfa*^*-/-*^ (IMSR_JAX:005540), *Adipoq*^*Cre/+*^ (IMSR_JAX:010803)^21^, *Atgl*^fl/fl^ (IMSR_JAX:024278)^22^. All experimental C57BL/6J WT, *Rag2*^*-/-*^, *Tnfa*^*-/-*^ mice were males between 8–12 weeks old. Sex and aged matched *Adipoq*^*Cre/+*^*Atgl*^*fl/fl*^ were used aged between 8-20 weeks old. Mice were housed in a Specific-Pathogen-Free barrier facility, at iMM, under standard laboratory conditions: 21 to 22°C ambient temperature and a 12h light/12h dark cycle. Chow and water were available ad libitum.

### Infection

*T. brucei* cryostabilates were thawed and parasite viability by its motility was confirmed under an optic microscope. Mice were infected by intraperitoneal (i.p.) injection of 2000 *T. brucei* parasites. At selected time-points post-infection, animals were euthanized by CO2 narcosis and immediately perfused transcardially with pre-warmed heparinised saline (50mL phosphate buffered saline (PBS) with 250 μL of 5000 I.U./mL heparin).

### Feeding synchronization

Feeding synchronization was achieved by fasting mice for 7 hours from 0 am to 7 am. Afterwards, access to food was re-established for 2 hours after which mice were euthanized.

### Paired feeding

Mice were individually housed, and their food intake monitored daily. To each infected mouse a non-infected control was assigned (i.e, pair-fed control). Every 24 hours, pair-fed controls were provided with the same amount of food consumed by their respective infected pair in the previous 24 hours.

### Body composition analysis

Mouse total fat mass, lean mass and free body fluid mass were determined using a 6.2 MHz time-domain nuclear magnetic resonance based Minispec LF65 (Bruker) apparatus. Live unanesthetized were restrained, weighed, and then inserted into the Minispec apparatus. Each measurement lasted for approximately one minute.

### Chemical sympathectomy

Chemical sympathectomy was performed by i.p. administration of 200mg/kg of 6-hydroxydopamine hydrobromide (6-OHDA) in PBS containing 0.4% ascorbic acid as stabilizer. 72 and 24 hours prior to infection, mice were treated with either 6-OHDA or with PBS containing 0.4% ascorbic acid (sham controls)^41^.

### Parasite lines

Mouse and in vitro infections were performed using parasites derived from T. brucei AnTat 1.1E, a pleomorphic clone derived from the EATRO1125 strain. AnTat 1.1E 90– 13 is a transgenic cell-line encoding the tetracyclin repressor and T7 RNA polymerase189. AnTat1.1E 90–13 GFP::PAD1utr derives from AnTat1.1E 90–13 in which the green fluorescent protein (GFP) is coupled to PAD1 3’UTR. Monomorphic bloodstream form Lister 427 parasites (MITat1.2, clone 221a) were used for in vitro infections. All parasite cell lines were propagated and maintained in HMI-11 medium at 37°C in a 5% CO2 atmosphere^42^.

### Ex vivo lipolysis assay

Follow euthanasia and systemic re-perfusion as previously described, AT depots were collected, rinsed with PBS and kept at 37°C in low-glucose (1 g/L) DMEM (Gibco, Thermo Fisher Scientific Inc.) without serum until processing. Next, AT depots were cut into ≈ 20 mg explants and incubated for 2 hours at 37°C in 96-well plates containing 200 μL low glucose DMEM with 5% (w/v) fatty acid-free BSA and 5 μM of Triacsin C (Sigma, T4540) per well. Up to 5 explants were used per AT depot. Next, free fatty acid (C8 and longer) and glycerol concentrations were assessed using commercially available colorimetric kits (MAK044 and MAK117, Sigma) according to the manufacturer’s instructions. Concentration of fatty acids and glycerol was normalized to each explant’s protein content. Protein contents were quantified by first delipidating the explants for 1 hour in 1 mL of 2:1 chloroform-methanol and 1% acetic acid solution at 37°C under vigorous agitation on a benchtop thermomixer. Afterwards, delipidated explants were lysed overnight in 500 μL of a 0.3M NaOH and 0.1% (w/v) SDS solution at 56°C under vigorous agitation. Lysate protein quantification was performed using a BCA Protein Assay Kit (A53225, Thermo Fisher Scientific Inc.) according to the manufacturer’s instructions. *Ex vivo* lipolysis rates are expressed as: Nanomoles of metabolite released per mg or protein per hour.

### Adipocyte lipid droplet area quantification

Images of hematoxylin and eosin-stained slides of gonadal AT were acquired in a Nanozoomer-SQ (Hamamatsu Photonics) with a magnification of 20x. Afterwards, at least 5 random fields per slide were analysed with imageJ using the Adiposoft (version 1.16) plugin^43^ using an adipocyte diameter range from 10 to 100 μm.

### Parasite quantification in blood and organs

For parasitemia quantification, blood samples were taken daily from the tail vein and diluted 1:150 in an PBS solution containing 2% formaldehyde. Parasites were counted manually in disposable 0.1 mm depth counting chambers (Kova International). Parasitemia detection limit is 3.75×10^5^ parasites per mL of blood (equivalent to one parasite counted in a total of 4 square with 0.1μL each).

For parasite quantification in organs, genomic DNA (gDNA) was extracted using NZY tissue gDNA isolation kit (NZYTech, Portugal). The amount of *T. brucei* 18S rDNA was measured by quantitative PCR (qPCR), using the primers 5’-ACGGAATGGCACCACAAGAC–3’ and 5’–GTCCGTTGACGGAATCAACC–3’, and converted into number of parasites using a calibration curve, as previously described^5^. Number of parasites per mg of organ (parasite density) was calculated by dividing the number of parasites by the mass of organ used for qPCR. The total amount of parasites in the organ was estimated by multiplying parasite density by the total mass of the organ.

### Preparation of single cell suspensions

Single cell suspensions containing AnTat1.1E 90–13 GFP::PAD1utr parasites were prepared for analysis of PAD1 expression by flow cytometry. Parasite adipose tissue forms were recovered by incubating the gonadal AT of infected mice in 5 mL of HMI-11 medium within 50 mL conical tubes under gentle agitation at 37°C for 30 min. Afterwards, cell suspensions were centrifuged at 2000 rpm, washed with PBS, fixed with 2% formaldehyde for 20 min, washed with PBS.

### Flow cytometry

Cell suspensions containing formaldehyde-fixed AnTat1.1E 90–13 GFP::PAD1utr parasites were stained with a PBS solution containing 0.005 mg/mL Hoechst 33342 (Thermo Fisher Scientific) for 20 min at 4°C.

Samples were passed through a 40μm-pore-size nylon cell strainer (BD Biosciences) and then analysed on a BD LSRFortessa flow cytometer with FACSDiva 6.2 Software. All data were analysed using FlowJo software version 10.0.7r2. Schematics of the gating strategies used parasite analysis are represented in **Supplementary Figure 5**.

### Statistical analysis

Statistical analysis was performed using GraphPad Prism (version 8.4.3). Data are presented as individual values or as mean ± SEM. Parasite numbers were transformed into their respective Log base 10 values to achieve linearization prior to statistical analysis. For analysis purposes, parasitemia values below the limit of detection were attributed a Log10 value equal to 5 (i.e. close the limit of detection). Statistical differences were assessed using two-way ANOVA and one-way ANOVA with Sidak’s test for multiple comparisons. Standalone comparisons were performed using an unpaired t test. P values lower than 0.05 were considered to be statistically significant.

## Acknowledgements

We are thankful to Bruno Silva-Santos and Marc Veldhoen for providing experimental mice. We are also grateful to the staff of iMM’s rodent facility for excellent animal husbandry and welfare services and to the staff of iMM’s Comparative Pathology Laboratory for expert technical assistance. This work was supported by European Research Council (771714), Fundação para a Ciência e Tecnologia (PD/BD/128286/2017 to H.M., CEECINST/00110/2018 to L.M.F.), by the Austrian Fonds zur Förderung der Wissenschaftlichen Forschung (F73 SFB Lipid Hydrolysis to R.Z.) and the Louis Jeantet Prize 2015 by the Fondation Louis Jeantet to R.Z.

## Supplementary figures

**Supplementary figure 1.**
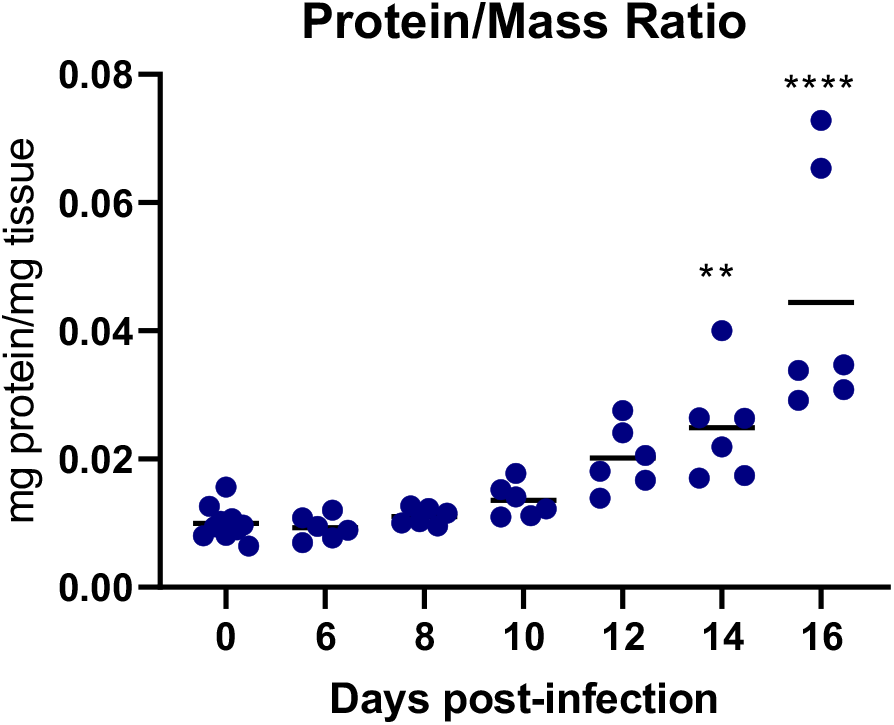
Dynamics of gonadal AT mass to protein ratio. Gonadal AT explant protein to total mass ratio. n= 5-10 mice per group. Statistical analysis was performed with One-way using Sidak’s test for multiple comparisons. *, P<0.05; **, P<0.01; ***, P<0.001; ****, P<0.0001.

**Supplementary figure 2.**
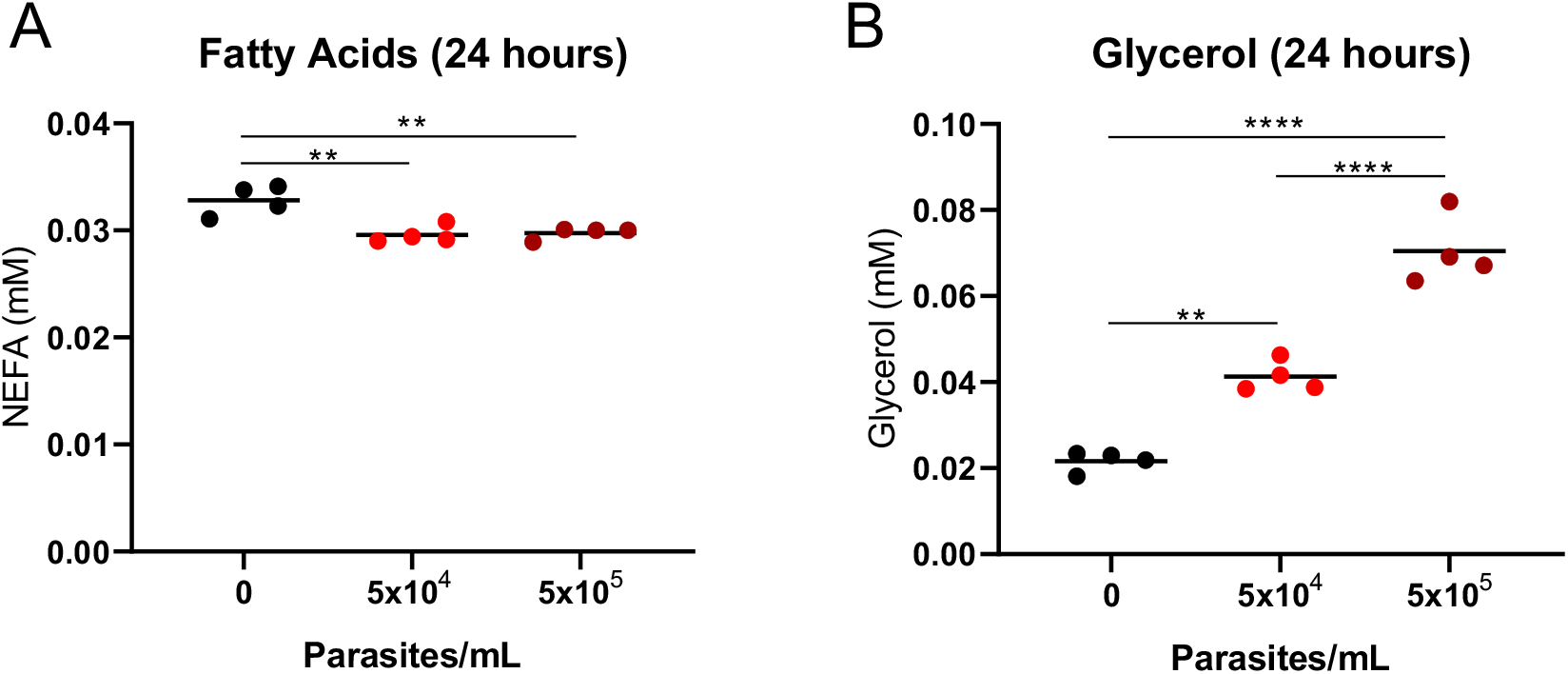
Parasite secretion of lipolytic metabolites. Concentration of **(A)** fatty acids and **(B)** glycerol in axenic HMI-11 cultures with 5% (w/v) BSA after 24 hours of incubation without parasites or with an initial inoculum of 5×10^4^ or 5×10^5^ parasites per mL. Statistical analysis was performed with one-way ANOVA using Sidak’s test for multiple comparisons. *, P<0.05; **, P<0.01; ***, P<0.001; ****, P<0.0001. Each data point represents an independent culture.

**Supplementary figure 3.**
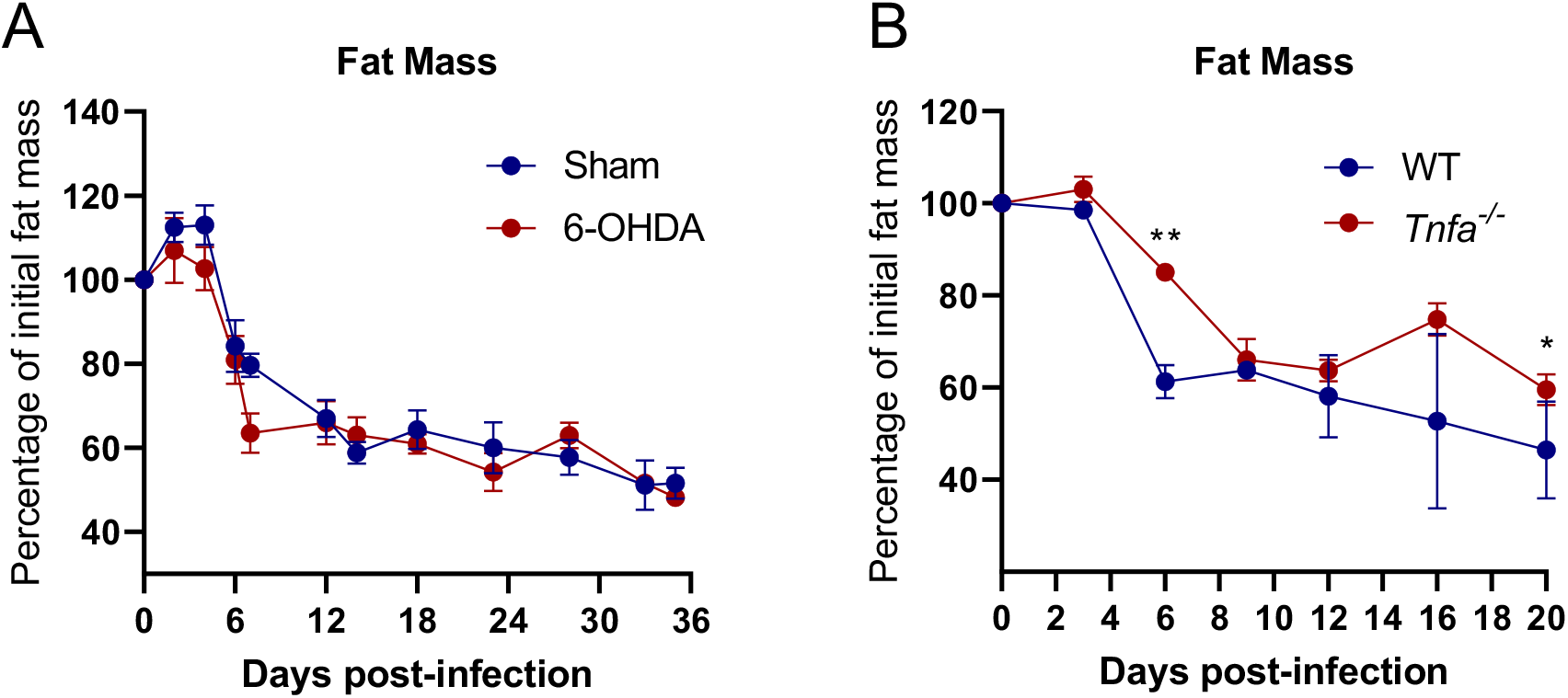
Effect of chemical sympathectomy and TNF-α deficiency on fat mass progression. Fat mass relative to baseline of infected **(A)** 6-OHDA-treated or **(B)** *Tnfa*^*-/-*^ mice and respective WT or sham-treated controls. Error bars represent the SEM (**(A)** n= 5 mice per group and **(B)** n= 3-5 mice per group). Statistical analysis was performed with a mixed-effects Two-way ANOVA using Sidak’s test for multiple comparisons. *, P<0.05; **, P<0.01; ***, P<0.001; ****, P<0.0001.

**Supplementary figure 4.**
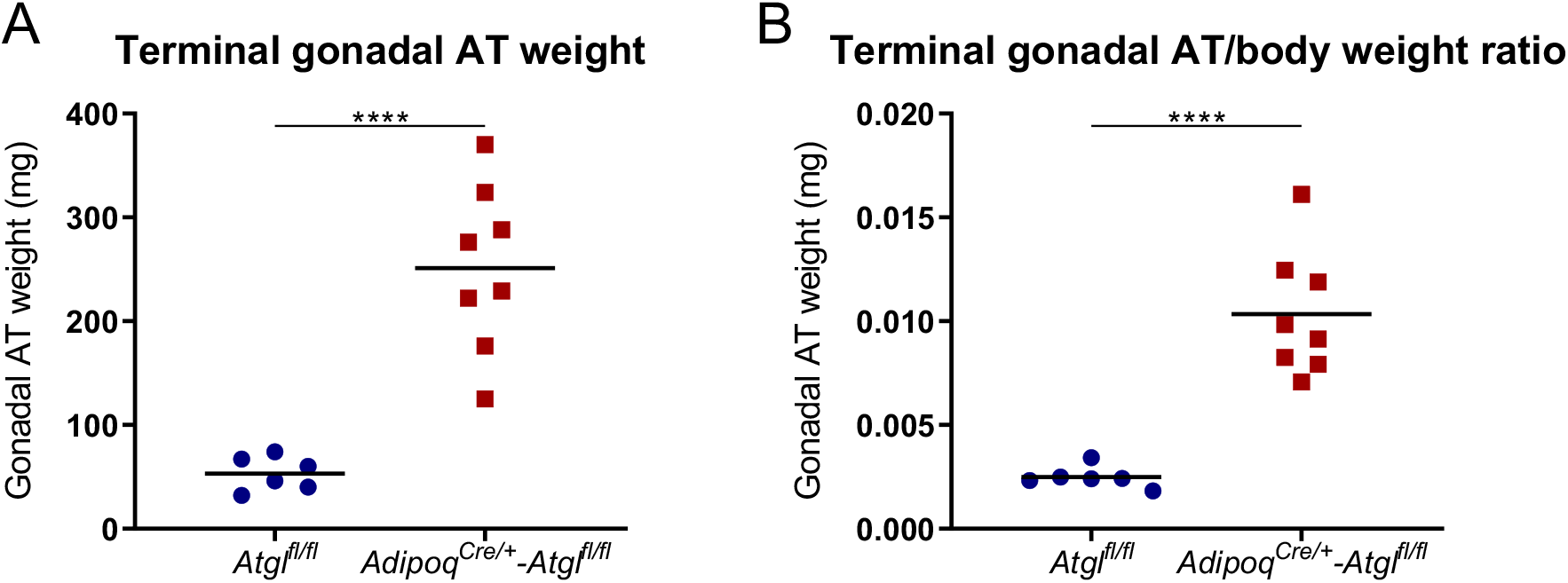
Effect of adipocyte-ATGL deficiency on terminal gonadal AT weight. Terminal **(A)** gonadal AT mass and **(B)** gonadal AT mass to body weight ratio of Adipoq^Cre/+^-*Atgl*^fl/fl^ and *Atgl*^fl/fl^ littermate controls. Error bars represent the SEM (**(A-B)** n=6-8 mice per group). Statistical analysis was performed with unpaired t test. *, P<0.05; **, P<0.01; ***, P<0.001; ****, P<0.0001.

**Supplementary figure 5.**
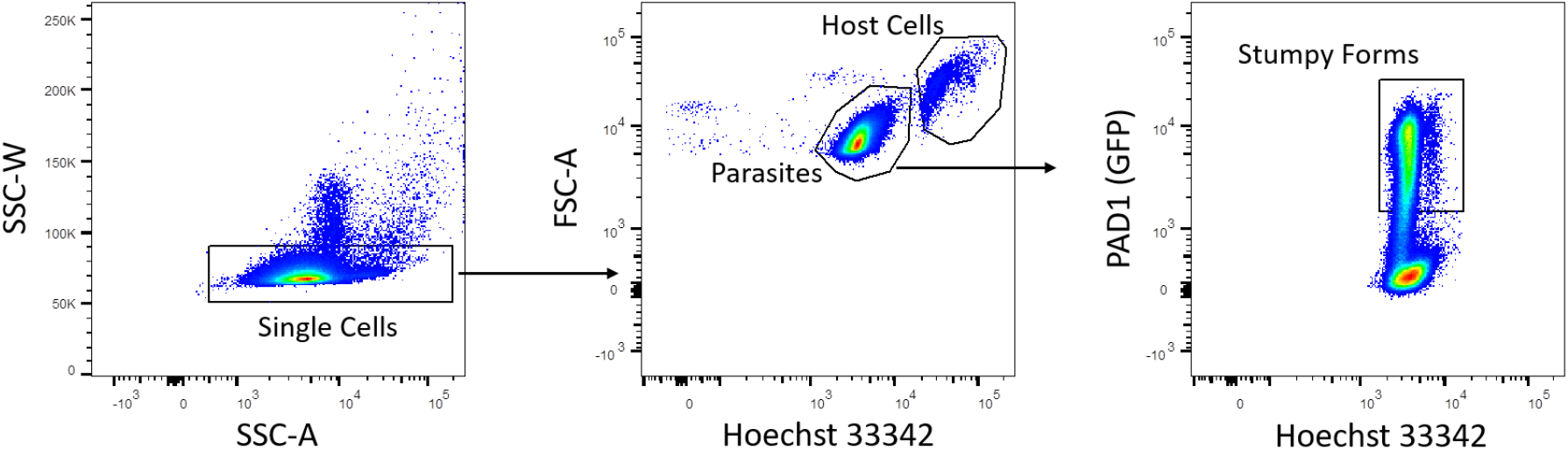
Parasite flow cytometry gating strategy. Single cells were identified using SSC-W vs SSC-A gating. Parasites and host cells were then differentiated based on Hoechst 33342 intensity. Stumpy forms of the parasite were identified based on PAD1 (GFP) expression.

## References

1. Franco, J. R., Simarro, P. P., Diarra, A. & Jannin, J. G. Epidemiology of human African trypanosomiasis. Clin. Epidemiol. 6, 257–275 (2014).

2. Pereira, S. S., Trindade, S., De Niz, M. & Figueiredo, L. M. Tissue tropism in parasitic diseases. Open Biol. 9, (2019).

3. Kennedy, P. G. E. Clinical features, diagnosis, and treatment of human African trypanosomiasis (sleeping sickness). Lancet. Neurol. 12, 186–94 (2013).

4. Camara, M. et al. Extravascular Dermal Trypanosomes in Suspected and Confirmed Cases of gambiense Human African Trypanosomiasis. Clin. Infect. Dis. 73, 12–20 (2021).

5. Trindade, S. et al. Trypanosoma brucei Parasites Occupy and Functionally Adapt to the Adipose Tissue in Mice. Cell Host Microbe 19, 837–848 (2016).

6. Machado, H. et al. Trypanosoma brucei triggers a broad immune response in the adipose tissue. PLoS Pathog. 17, 1–26 (2021).

7. Grabner, G. F., Xie, H., Schweiger, M. & Zechner, R. Lipolysis: cellular mechanisms for lipid mobilization from fat stores. Nat. Metab. 3, 1445–1465 (2021).

8. Gasic, S., Tian, B. & Green, A. Tumor necrosis factor α stimulates lipolysis in adipocytes by decreasing G(i) protein concentrations. J. Biol. Chem. 274, 6770–6775 (1999).

9. White, J. E. & Engel, F. L. A Lipolytic Action of Epinephrine and Norepinephrine on Rat Adipose Tissue in vitro. Proc. Soc. Exp. Biol. Med. 99, 375–378 (1958).

10. Chakrabarti, P. et al. Insulin Inhibits Lipolysis in Adipocytes via the Evolutionarily Conserved mTORC1-Egr1-ATGL-Mediated Pathway. Mol. Cell. Biol. 33, 3659–3666 (2013).

11. Zu, L. et al. Bacterial endotoxin stimulates adipose lipolysis via toll-like receptor 4 and extracellular signal-regulated kinase pathway. J. Biol. Chem. 284, 5915–5926 (2009).

12. Chi, W. et al. Bacterial peptidoglycan stimulates adipocyte lipolysis via NOD1. PLoS One 9, 1–9 (2014).

13. Rosen, E. D. & Spiegelman, B. M. Adipocytes as regulators of energy balance and glucose homeostasis. Nature 444, 847–853 (2006).

14. Das, S. K. et al. Adipose Triglyceride Lipase Contributes to Cancer-Associated Cachexia. Science (80-.). 333, 233–238 (2011).

15. Morigny, P., Houssier, M., Mouisel, E. & Langin, D. Adipocyte lipolysis and insulin resistance. Biochimie 125, 259–266 (2016).

16. Geng, Y., Faber, K. N., de Meijer, V. E., Blokzijl, H. & Moshage, H. How does hepatic lipid accumulation lead to lipotoxicity in non-alcoholic fatty liver disease? Hepatol. Int. 15, 21–35 (2021).

17. Eisenthal, R. & Panes, A. The aerobic/anaerobic transition of glucose metabolism in Trypanosoma brucei. FEBS Lett. 181, 23–27 (1985).

18. Michels, P. A. M., Bringaud, F., Herman, M. & Hannaert, V. Metabolic functions of glycosomes in trypanosomatids. Biochim. Biophys. Acta - Mol. Cell Res. 1763, 1463–1477 (2006).

19. Malmfors, T. & Sachs, C. Degeneration of adrenergic nerves produced by 6-hydroxydopamine. Eur. J. Pharmacol. 3, 89–92 (1968).

20. Aresta-Branco, F., Sanches-Vaz, M., Bento, F., Rodrigues, J. A. & Figueiredo, L. M. African trypanosomes expressing multiple VSGs are rapidly eliminated by the host immune system. Proc. Natl. Acad. Sci. U. S. A. 116, 20725–20735 (2019).

21. Eguchi, J. et al. Transcriptional Control of Adipose Lipid Handling by IRF4. Cell Metab. 13, 249–259 (2011).

22. Sitnick, M. T. et al. Skeletal Muscle Triacylglycerol Hydrolysis Does Not Influence Metabolic Complications of Obesity. Diabetes 62, 3350–3361 (2013).

23. Morrison, L. J. et al. A major genetic locus in Trypanosoma brucei is a determinant of host pathology. PLoS Negl. Trop. Dis. 3, 1–12 (2009).

24. De Niz, M. et al. Organotypic endothelial adhesion molecules are key for Trypanosoma brucei tropism and virulence. Cell Rep. 36, (2021).

25. Matthews, K. R. Trypanosome Signaling — Quorum Sensing. (2021).

26. Kaur, S. et al. Adipose-specific ATGL ablation reduces burn injury-induced metabolic derangements in mice. Clin. Transl. Med. 11, (2021).

27. Baazim, H. et al. CD8+ T cells induce cachexia during chronic viral infection. Nat. Immunol. 20, 701–710 (2019).

28. Rouzer, C. A. & Cerami, A. Hypertriglyceridemia associated with Trypanosoma brucei brucei infection in rabbits: Role of defective triglyceride removal. Mol. Biochem. Parasitol. 2, 31–38 (1980).

29. Ebadi, M. & Mazurak, V. C. Evidence and mechanisms of fat depletion in cancer. Nutrients 6, 5280–5297 (2014).

30. Petruzzelli, M. & Wagner, E. F. Mechanisms of metabolic dysfunction in cancer-associated cachexia. Genes Dev. 30, 489–501 (2016).

31. Tracey, K. J. & Cerami, A. Metabolic responses to cachectin/TNF. A brief review. Ann. N. Y. Acad. Sci. 587, 325–331 (1990).

32. Baazim, H., Antonio-Herrera, L. & Bergthaler, A. The interplay of immunology and cachexia in infection and cancer. Nat. Rev. Immunol. 22, 309–321 (2022).

33. Delano, M. J. & Moldawer, L. L. The oriqins of cachexia in acute and chronic inflammatory diseases. Nutr. Clin. Pract. 21, 68–81 (2006).

34. Lafontan, M. & Langin, D. Lipolysis and lipid mobilization in human adipose tissue. Prog. Lipid Res. 48, 275–297 (2009).

35. Zechner, R., Madeo, F. & Kratky, D. Cytosolic lipolysis and lipophagy: Two sides of the same coin. Nat. Rev. Mol. Cell Biol. 18, 671–684 (2017).

36. Sinton, M. C. et al. Interleukin-17 drives sex-dependent weight loss and changes in feeding behaviour during Trypanosoma brucei infection. bioRxiv 2022.09.23.509158 (2022).

37. Grant, R. W. & Stephens, J. M. Fat in flames: Influence of cytokines and pattern recognition receptors on adipocyte lipolysis. Am. J. Physiol. - Endocrinol. Metab. 309, E205–E213 (2015).

38. Rojas, F. et al. Oligopeptide Signaling through TbGPR89 Drives Trypanosome Quorum Sensing. Cell 176, 306–317.e16 (2019).

39. Nolan, S. J., Romano, J. D., Kline, J. T. & Coppens, I. Novel approaches to kill toxoplasma gondii by exploiting the uncontrolled uptake of unsaturated fatty acids and vulnerability to lipid storage inhibition of the parasite. Antimicrob. Agents Chemother. 62, (2018).

40. Dawoody Nejad, L., Serricchio, M., Jelk, J., Hemphill, A. & Bütikofer, P. TbLpn, a key enzyme in lipid droplet formation and phospholipid metabolism, is essential for mitochondrial integrity and growth of Trypanosoma brucei. Mol. Microbiol. 109, 105–120 (2018).

41. Cardoso, F. et al. Neuro-mesenchymal units control ILC2 and obesity via a brain–adipose circuit. Nature 597, 410–414 (2021).

42. Hirumi, H. & Hirumi, K. Continuous cultivation of Trypanosoma brucei blood stream forms in a medium containing a low concentration of serum protein without feeder cell layers. J. Parasitol. 75, 985–989 (1989).

43. Galarraga, M. et al. Adiposoft: Automated software for the analysis of white adipose tissue cellularity in histological sections. J. Lipid Res. 53, 2791–2796 (2012).

